# Modulatory potentials of zerumbone isolated from ginger (*Zingiber zerumbet*) on eicosanoids: evidence from LPS induced peripheral blood leukocytes

**DOI:** 10.1101/2021.10.29.465185

**Authors:** Vinayak Uppin, Hamsavi Gopal Kamala, Bettadaiah Bheemanakere Kempaiah, Ramaprasad Ravichandra Talahalli

## Abstract

Several bioactive molecules from plant origin have been studied for their anti-inflammatory properties. In this study, we deciphered the anti-eicosanoid properties of zerumbone (sesquiterpene) isolated from ginger (*Zingiber zerumbet*) in LPS induced peripheral blood leukocytes from rats. Molecular interaction between zerumbone (Z) and eicosanoid metabolizing enzymes (COX-2, 5-LOX, FLAP, and LTA_4_-hydrolase) and receptors (EP-4, BLT-1, and ICAM-1) along with NOS-2 were assessed using Auto-Dock 4.2 docking software. Further, the rat peripheral blood leukocytes were isolated and treated with zerumbone (5μM) and activated using bacterial lipopolysaccharide (10nM). Oxidative stress (OS) markers, reactive oxygen species, antioxidant enzymes, COX-2, 5-LOX, BLT-1, EP-4 were assessed along with the activity of COX-2. Zerumbone showed a higher binding affinity with mPGES-1, NOS-2, FLAP, COX-2, LTA-4-hydrolase, and BLT-1 mediators of the eicosanoid pathway. Further, zerumbone significantly (p<0.05) inhibited COX-2, 5-LOX, NOS-2, EP-4, BLT-1, and ICAM-1 expression in LPS induced peripheral blood leukocytes from rats. Zerumbone positively modulates critical enzymes and receptors of eicosanoids in leukocytes activated with lipopolysaccharides. Thus, zerumbone offers a promising therapeutic strategy in the management of inflammation.

## Introduction

Zerumbone (2, 6, 9, 9-tetramethyl-[2*E*, 6E, 10E]-cycloundeca-2, 6, 10-trien-1-one) is a monocyclic sesquiterpene present in Zingiberaceae family. Chemically, zerumbone has three double bonds with a conjugated carbonyl group in an 11-membered ring structure [1]. Many investigators using cell lines and animal models have addressed the anti-cancerous [2], hepatoprotective [3], hypolipidemic [4], antioxidative [5], and neuroprotective [6], effects of zerumbone (Z). Our previous studies revealed that zerumbone prevents cognition loss induced by hyperlipidemia in rats [7]. In the presence of long-chain n-3 fatty acids (EPA, 20:5n-3 + DHA, 22:6n-3), zerumbone presented greater effects in preventing hyperlipidemia-induced cognitive decline [7]. Because of the presence of α, β-unsaturated carbonyl-based moiety in its structure [8], zerumbone may potentially alter the inflammatory response triggered in the body. Studies have demonstrated that zerumbone weakens nuclear factor kappa-B (NF-kB) signaling and nitric oxide production to cause the resolution of inflammation [9]. However, there are no reports on the modulatory effects of zerumbone on eicosanoid metabolism. Prostaglandins and leukotrienes (collectively termed eicosanoids) are potent lipid mediators of inflammation derived by cyclooxygenase and 5-lipoxygenase, respectively [10]. Eicosanoids may amplify or reduce inflammation, which coordinates cytokine production, antibody formation, cell proliferation and migration, and antigen presentation [11]. The eicosanoids act through specific G protein-coupled receptors and can be regulated by nonsteroidal anti-inflammatory drugs. Studies with knockout mice and chimeras have established the implications of eicosanoids in general and leukocytes, particularly in the pathogenesis of inflammatory complications [12, 13]. The central role of circulating leukocytes in the initiation, propagation, and resolution of inflammation is well established by other investigators [14]. In this study, we deciphered zerumbone’s anti-inflammatory role mediated through the downregulation of eicosanoids in LPS treated rat peripheral blood leukocytes. For the first time, our finding supports the therapeutic application of zerumbone in managing inflammation via countering the eicosanoids.

## Materials and methods

### Materials

Acrylamide, ammonium per-sulfate, adenosine diphosphate, BSA, beta-mercaptoethanol, CDNB, cytochrome-C, glycine, thiobarbituric acid, Tris-HCl, xanthine, LPS (L4391), and xanthine oxidase were obtained from Sigma Chemicals, St. Louis, MO, USA. Dinitrophenyl hydrazine, EDTA, glutathione oxidized, glutathione reduced, malonaldehyde, NADPH, phosphoric acid, sulphanilamide, sulphosalicylic acid, sodium nitrate, t-butyl hydroperoxide were obtained from SRL Chemicals, Mumbai, India. The ELISA kit for measurement of COX-2 activity was purchased from Cayman Chemicals, Ann Arbor, MI, USA. Zerumbone (99.9% pure) was extracted from ginger as per the earlier procedure [15].

### Molecular docking study

The 3D structural coordinates of Zerumbone [CID-5470187] were retrieved from the Pubchem database [16], and ligand preparation was carried out using Auto-Dock Tools 4.2. The X-ray diffraction-based crystal structure of target proteins was downloaded from RCSB-PDB. Target proteins such as Microsomal Prostaglandin E Synthase-1, mPGES-1 [4yk5],, Inter cellular adhesion molecule-1, ICAM-1 [1mqs], Nitric oxide synthase-2, NOS-2 [1m9t], Cyclooxygenase-2, COX-2 [5ikv], Leukotriene B_4_ receptor-1, BLT-1 [5×33], 5-Lipoxygenase activating protein, FLAP [2q7r], Leukotriene A_4_ hydrolase, LTA4-H [5n3w], and Prostaglandin-E receptor-4, EP-4 [5ywy], in complex with inhibitors and resolution of <3.0Å were selected for the study. The proteins were prepared by removing water molecules, adding polar hydrogen’s, Kollman united atom charges, and Gasteiger charges to the ligand. The complexes bound to the protein receptor molecule were removed, and molecular docking simulation was carried out using Auto-Dock 4.2. Lamarckian genetic algorithm (LGA) method was followed to find the optimal conformation of the ligand. Grid box dimensions were kept minimal, covering the proteins’ active site residues, and grid maps were generated. Using LGA, molecular docking was carried out, and ten possible outcomes were generated, the one with the lowest binding energies picked up from the cluster. The visualization of the docked complex files was performed by UCSF chimera, and a 2D interaction map of the complex was generated using Lipgplot^+^ software.

### Isolation of peripheral blood leukocytes and treatment

The experimental animal procedures followed in this study were approved (approval number CFT/IAEC/120/2018) and carefully monitored by the Institutional Animal Care and Use Committee of CSIR-Central Food Technological Research Institute, Mysore, India. Whole blood from rats fed on a regular pellet diet was collected by cardiac puncture and transferred to vacutainer tubes containing K_2_ EDTA. Ice-cold RBC lysis buffer (4 volumes) was added to whole blood, mixed gently, and centrifuged at 2000rpm (291 RCF) for 15min. The supernatant was discarded, and the step was repeated to pellet the circulating leukocytes. The leukocyte pellet was suspended in Hank’s Balanced Salt Solution (HBSS) containing 1% fetal bovine serum (FBS).

#### Treatment

An equal number (5 x 10^6^) of leukocytes were treated with or without zerumbone (1μM, 2.5μM, 5μM, 10μM, and 20μM) for 60 min at 37^0^C and then activated with bacterial lipopolysaccharide (LPS, 10nM) for 2hrs. Control cells were neither incubated with zerumbone or LPS. The samples were centrifuged at 3000 rpm for 15 min, and the supernatant was taken for the evaluation of generation of reactive oxygen species and nitric oxide production. After determining the optimum concentration of zerumbone, further experiments were carried out at 5μM zerumbone concentration. Upon completion of the treatments, the leukocytes were made into pellet by centrifugation at 2000 rpm (291 RCF), and was lysed using cell lysis buffer supplied by cell signalling, USA. The protein content in the cell lysate was measured by Lowry’s method [17].

### Measurement of reactive oxygen species, oxidative stress markers, and antioxidant enzymes

The production of reactive oxygen species (ROS) in leukocytes was measured using Dichlorofluorescein diacetate (DCF-DA) and the fluorescent product 2, 7, dichlorofluorescein DCF at the excitation wavelength of 488nm and emission at 525nm [18]. The oxidative stress (OS) markers and antioxidant enzymes activity was measured using cell lysate prepared using phosphate buffer saline (PBS) buffer. The aliquots of supernatants were taken for the measurement of lipid peroxides [19], Nitric oxide [20] and protein carbonyls [21]. The activity of antioxidant defense enzymes, including catalase [22], SOD [23], glutathione peroxidase [24], and glutathione transferase [25] were measured spectrophotometrically.

### Western immunoblotting and ELISA

Equal amount of protein was loaded and resolved on 10% polyacrylamide gel and transferred onto the PVDF membrane supplied by Bio-Rad, USA. The membranes were blocked with 5% non-fat dry milk and probed for ICAM-1, COX-2, and GAPDH (Cloud-Clone Corp., TX, USA), NOS-2, EP-4, (Santa Cruz Biotechnology, CA, USA), BLT-1, 5-LOX, (Abcam, Cambridge, UK) overnight at 4°C. The membranes after three washes with TBST were incubated with appropriate secondary antibodies. Upon completion of 3 washes with TBST and one wash with TBS, the membranes were treated with enhanced chemiluminescent reagents obtained from Bio-Rad (Bio-Rad Laboratories. Inc., USA) and documented by Syngene Gel Doc (G: BOX Chemi XT4) for densitometry.

The COX-2 activity was assayed as per the kit instructions [400012] supplied by the Cayman-Chemicals (Ann Arbor, MI, USA). The colorimetric determination of the oxidized compound N, N, N, N-Tetramethyl-p-phenylenediamine (TMPD) at 590 nm was determined spectrophotometrically. The eicosanoid level in the peripheral blood leukocytes preparation was measured by the ELISA kits supplied by Cayman Chemicals (Ann Arbor, MI, USA). Briefly, 1 x 10^6^ million cells were suspended in the Hank’s Balanced Salt Solution (HBSS) containing 0.1% fetal bovine serum (FBS). To assess the inhibitory properties, the cells pre-incubated with zerumbone (5uM) for 1 hr, followed by LPS (10nM) for 2 hrs. The treated cells were separated by centrifugation (10,000rpm), and the supernatant was used for the measurement of eicosanoids.

### Statistical analysis

The sample size was determined based on the studies reported earlier, with a statistical power of 0.8 and α error of 0.05. The data were analyzed by One-way ANOVA (non-parametric), followed by Tukey’s test using Graph Pad Prism version 7.04, and a P-value < 0.05 was considered statistically significant. The false discovery rate (q value of 0.1) was considered as the threshold for significance. Only the mean and standard deviation for each diet group are plotted in each figure.

## Results

### Molecular interaction between zerumbone and target proteins

Molecular docking reveals an appropriate ligand that fits both energetically and geometrically to the protein’s binding site. Among the proteins evaluated for the binding affinity with zerumbone, mPGES-1, NOS-2, FLAP, COX-2, LTA_4_-H, and BLT-1 showed the lowest binding energy (Kcal/mol) and lowest dissociation constant, suggesting the highest binding affinity. The binding affinity, dissociation constant (Ki), and amino acid residues involved in the interactions are given in Table-1. Fig. 1 depicts the binding pocket site and the hydrophobic interaction map of zerumbone.

**Figure 1.**
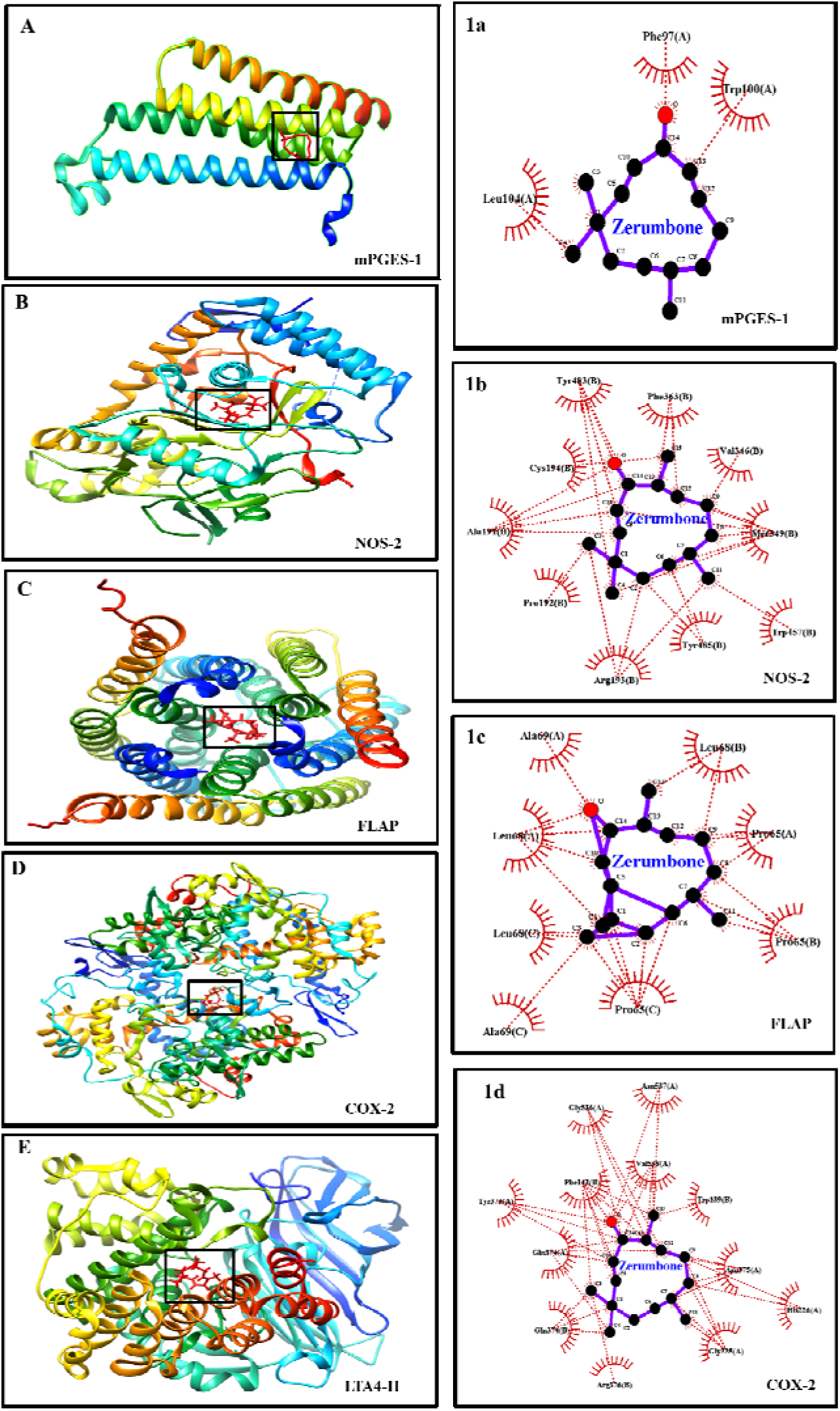
Interaction of zerumbone with target proteins including mPGES-1 (A), NOS-2 (B), FLAP (C), COX-2 (D), and LTA_4_-H (E). 1a, 1b, 1c, and 1d are the hydrophobic interaction of the zerumbone with amino acid residues of (A), (B), (C) and (D). The molecular docking was done by using Autock Tools 4.2 software and visualization was carried out by UCSF chimera and Lig-Plot^+^ softwares.

**Table 1.**
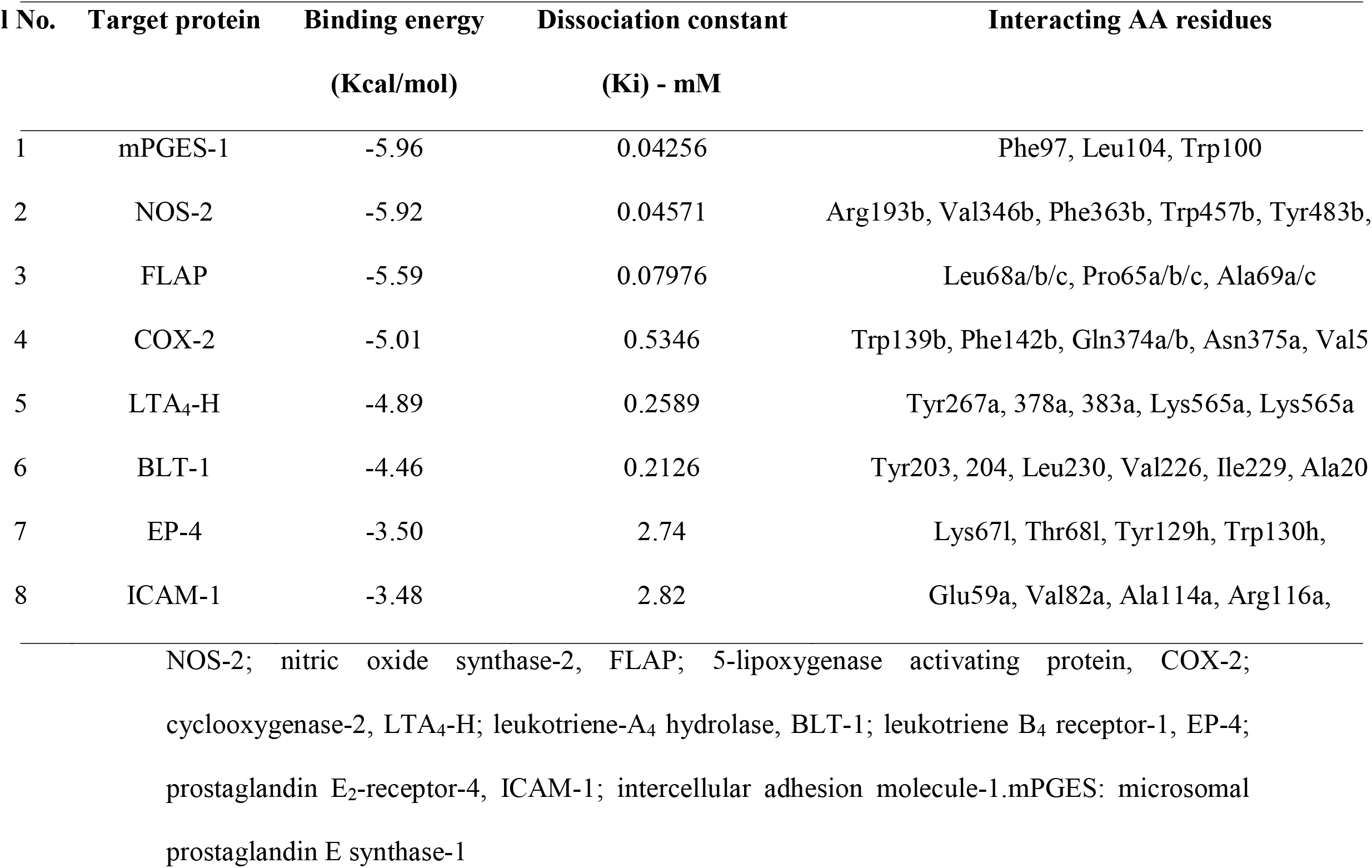
Binding affinity of zerumbone with target proteins

### Dose dependent effect of zerumbone on generation of ROS and NO release in peripheral blood leukocytes

The ROS level in leukocytes treated with LPS was significantly (p<0.05) increased compared to control. Whereas, the leukocyte pre-treatment with different doses of zerumbone (1 μM, 2.5 μM, 5 μM, 10 μM, and 20 μM) exhibited dose dependent decrease in the generation of ROS and NO production (Table 2). However, for further experiments a single concentration of 5μM zerumbone was used in order to elucidate the anti-eicosanoids effects of zerumbone in LPS induced leukocytes.

**Table.2.** Dose dependent effect of zerumbone on generation of ROS and NO release in peripheral blood leukocytes

| **Groups** | **Reactive oxygen species**<br>**(RFU/million cells)** | **Nitric oxide**<br>**(nmoles/mg protein)** |
| --- | --- | --- |
| **Control** | 16937.67 ± 762.05<sup>a</sup> | 28.54 ± 0.91<sup>a</sup> |
| **LPS** | 20229.01 ± 680.7<sup>b</sup> | 39.70 ± 1.67<sup>b</sup> |
| **LPS + Z (1μM)** | 19443.17 ± 1120.1<sup>a</sup> | 39.29 ± 2.28<sup>b</sup> |
| **LPS + Z (2.5μM)** | 18457.67 ± 157.5<sup>a</sup> | 37.41 ± 1.21<sup>a</sup> |
| **LPS + Z (5μM)** | 17650.15 ± 517.5<sup>a</sup> | 29.63 ± 3.35<sup>a</sup> |
| **LPS + Z (10μM)** | 17178.16 ± 206.03<sup>a</sup> | 26.47 ± 0.16<sup>a</sup> |
| **LPS + Z (20μM)** | 16650.08 ± 860.36<sup>a</sup> | 25.00 ± 2.05<sup>a</sup> |

Values not sharing a common superscript within the column are statistically significant at $p < 0.05$ . Values are Mean ± SD of 4 samples. The data were analyzed by ANOVA (non-parametric), followed by Tukey's test. LPS; Lipopolysaccharide, Z; zerumbone.

### Measurement of OS markers and antioxidant enzymes in peripheral blood leukocytes

The OS markers, including lipid peroxides and protein carbonyls, were significantly (p<0.05) increased in LPS treated leukocytes compared to control (Table-3). Pre-treatment with zerumbone significantly (p<0.05%) decreased the OS markers such as lipid peroxides, and protein carbonyls compared to LPS treated leukocytes. The activities of antioxidant enzymes, including catalase, SOD, glutathione peroxidase, and glutathione transferase, were significantly (p<0.05) decreased in LPS treated cells when compared to control (Table-3). Whereas pretreatment with zerumbone, significantly (p<0.05%), increased antioxidant enzymes’ activity compared to LPS treated leukocytes.

**Table 3.**
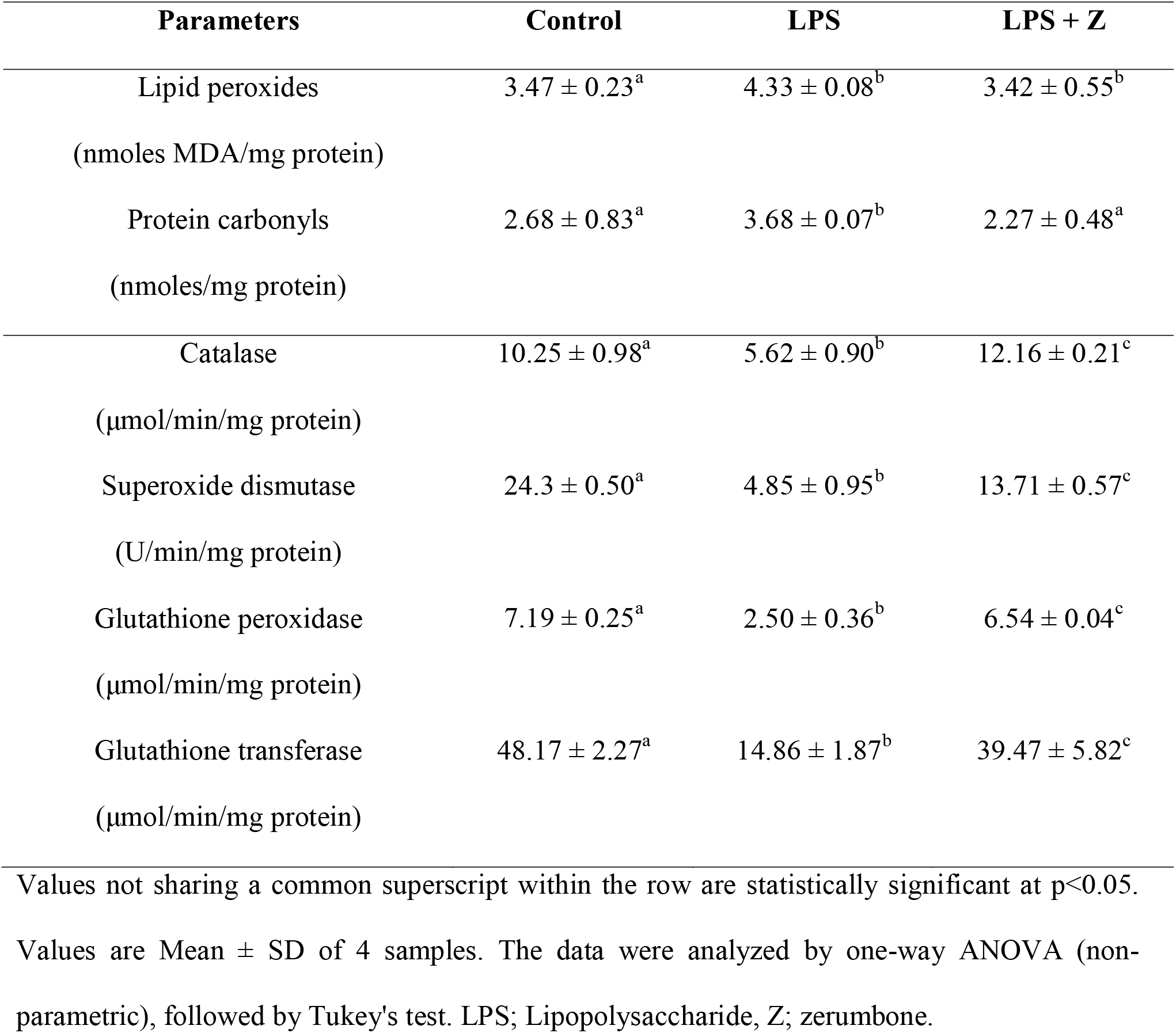
Oxidative stress markers and antioxidant enzyme activity in peripheral blood leukocytes.

| **Parameters** | **Control** | **LPS** | **LPS + Z** |
| --- | --- | --- | --- |
| Lipid peroxides<br>(nmoles MDA/mg protein) | $3.47 \pm 0.23^a$ | $4.33 \pm 0.08^b$ | $3.42 \pm 0.55^b$ |
| Protein carbonyls<br>(nmoles/mg protein) | $2.68 \pm 0.83^a$ | $3.68 \pm 0.07^b$ | $2.27 \pm 0.48^a$ |
| Catalase<br>($\mu\text{mol}/\text{min}/\text{mg}$ protein) | $10.25 \pm 0.98^a$ | $5.62 \pm 0.90^b$ | $12.16 \pm 0.21^c$ |
| Superoxide dismutase<br>(U/min/mg protein) | $24.3 \pm 0.50^a$ | $4.85 \pm 0.95^b$ | $13.71 \pm 0.57^c$ |
| Glutathione peroxidase<br>($\mu\text{mol}/\text{min}/\text{mg}$ protein) | $7.19 \pm 0.25^a$ | $2.50 \pm 0.36^b$ | $6.54 \pm 0.04^c$ |
| Glutathione transferase<br>($\mu\text{mol}/\text{min}/\text{mg}$ protein) | $48.17 \pm 2.27^a$ | $14.86 \pm 1.87^b$ | $39.47 \pm 5.82^c$ |

Values not sharing a common superscript within the row are statistically significant at $p < 0.05$ .
Values are Mean ± SD of 4 samples. The data were analyzed by one-way ANOVA (non-parametric), followed by Tukey's test. LPS; Lipopolysaccharide, Z; zerumbone.

### Expression of COX-2, EP-4, 5-LOX, BLT-1, and activity of COX-2 in peripheral blood leukocytes

The expression of COX-2, EP-4, 5-LOX, BLT-1 in leukocytes treated with LPS was significantly (p<0.05) increased when compared to control (Fig. 2). Pre-treatment with zerumbone significantly (p<0.05%) decreased COX-2, EP-4, 5-LOX, and BLT-1 expression compared to LPS treated leukocytes. Further, the activity of COX-2 in LPS treated leukocytes was significantly (p<0.05) higher when compared to control (Fig. 3). Whereas pre-treatment with zerumbone significantly decreased the activity of COX-2 when compared to LPS treated leukocytes.

**Figure 2.**
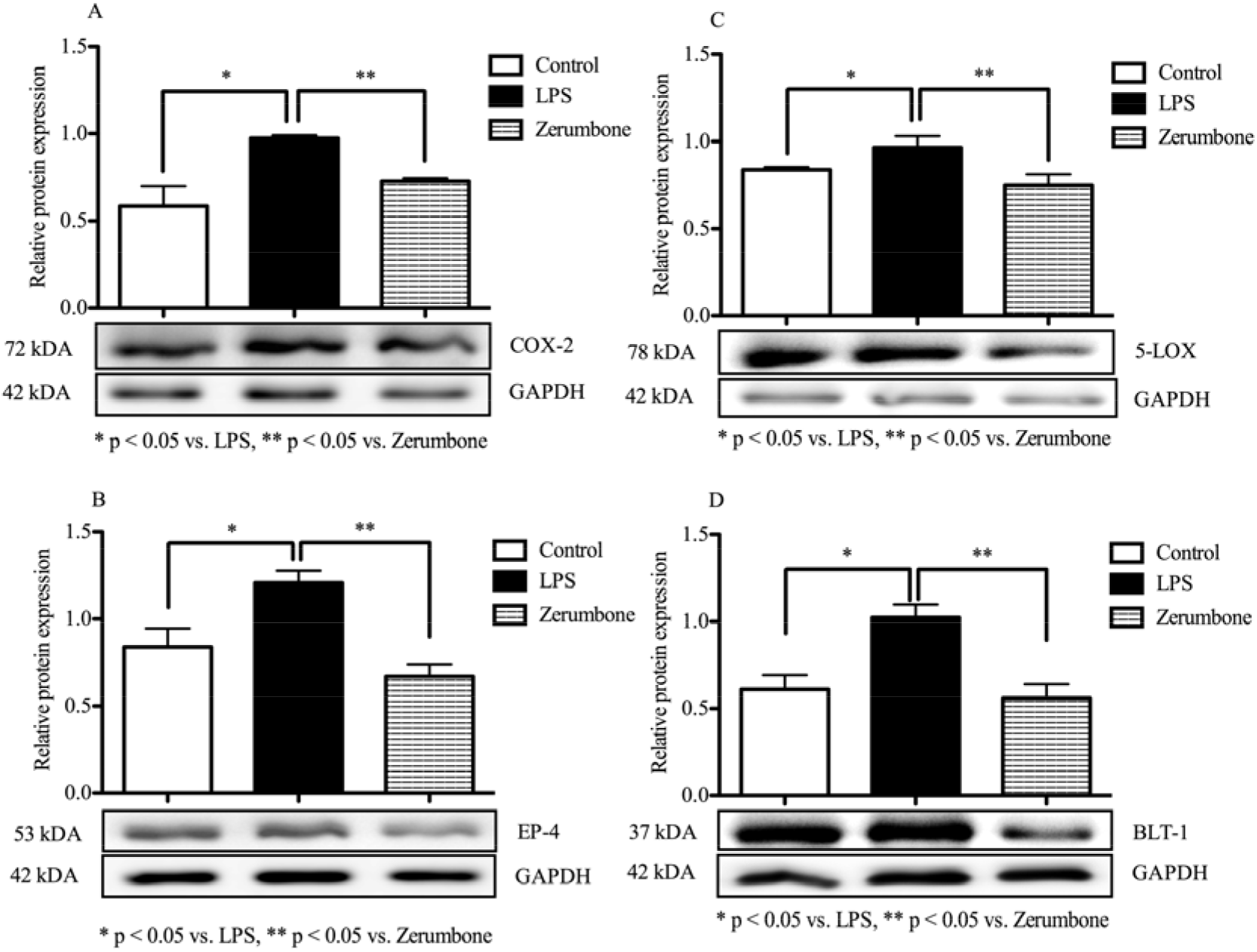
Relative expressions of COX-2 (A), EP-4 (B), 5-LOX (C), and BLT-1 (D) proteins in peripheral blood leukocytes treated with 5μM zerumbone. Values are Mean ± SD of 3 samples. A p value < 0.05 was considered statistically significant.

**Figure 3.**
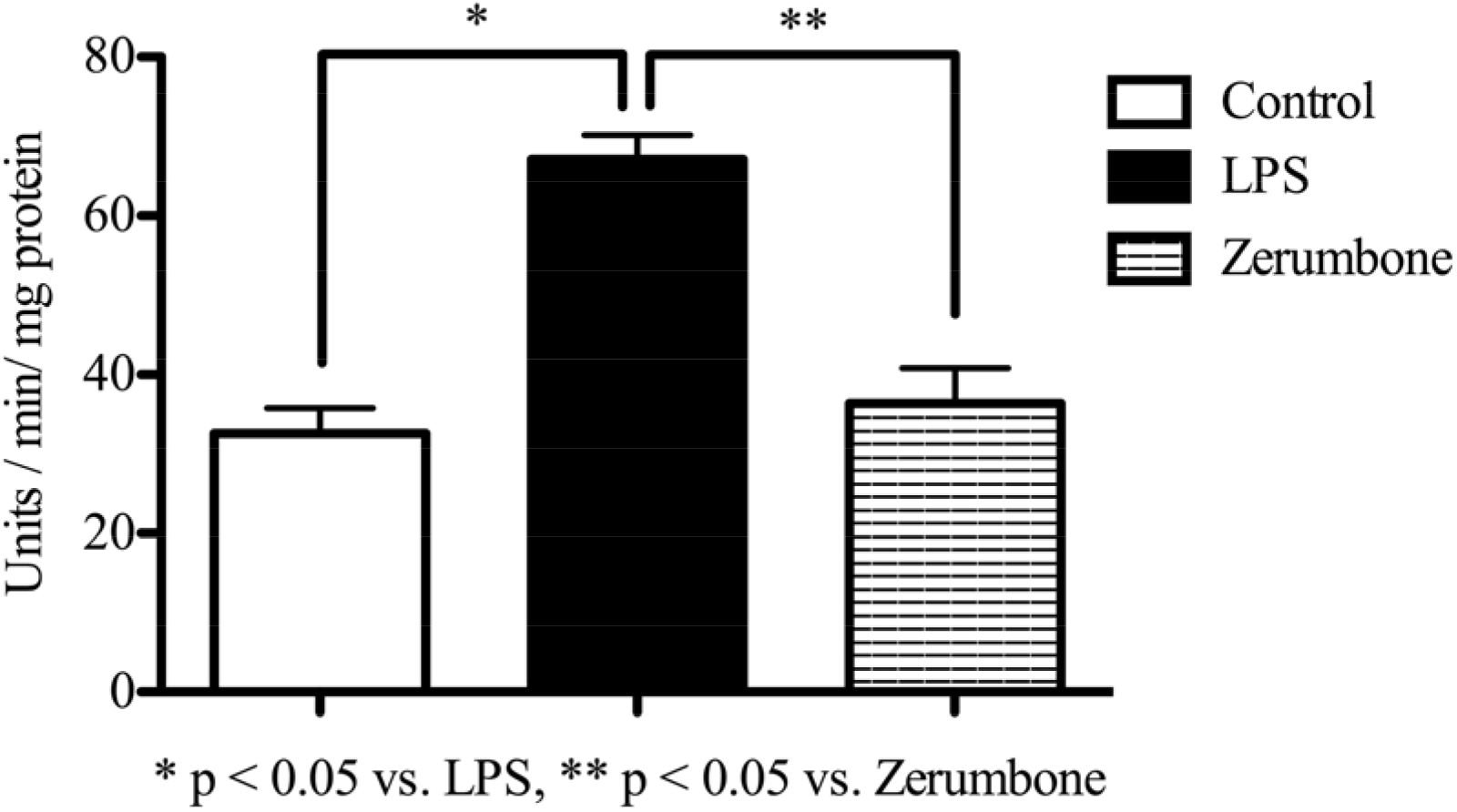
Activity of COX-2 in peripheral blood leukocytes treated with 5μM zerumbone. Values are Mean ± SD of 6 samples. A p value < 0.05 was considered statistically significant.

### Eicosanoid production in peripheral blood leukocytes

The levels of PGE2, LTB4, and LTC4 in leukocytes treated with LPS were found to be increased by 128, 179, and 151%, respectively, compared to control (Fig. 4). Whereas, pre-treatments with zerumbone decreased the LPS induced production of PGE_2_, LTB_4_, and LTC_4_ by 28, 45, and 48%, respectively, compared to the leukocytes stimulated without pre-treatment with zerumbone.

**Figure 4.**
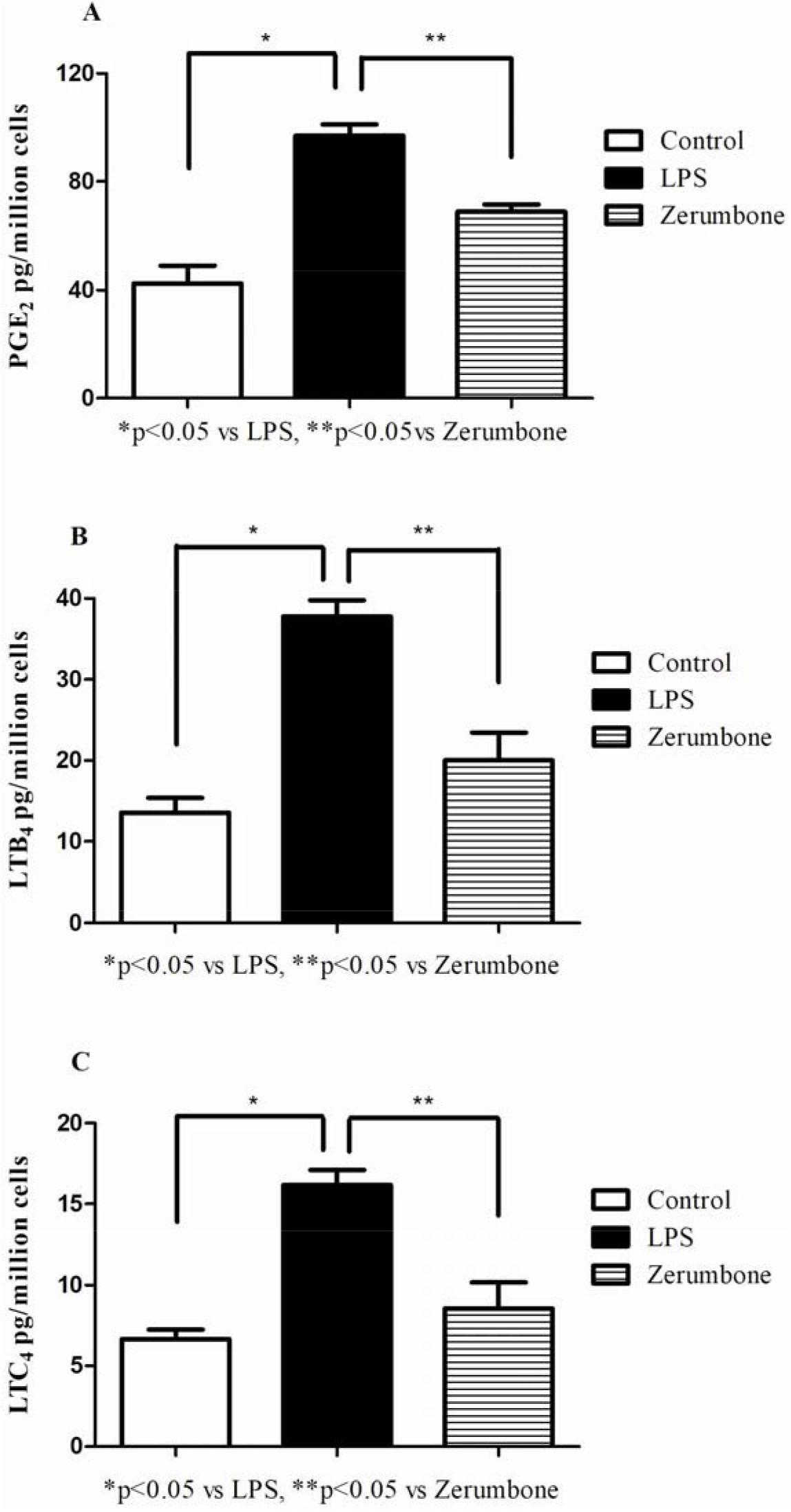
Generation of PGE_2_ (A), LTB_4_ (B), and LTC_4_ (C) in peripheral blood leukocytes Values are Mean ± SD of 3 samples. A p value < 0.05 was considered statistically significant.

### Expression of NOS-2, and ICAM-1 in peripheral blood leukocytes

The NOS-2 expression in the LPS treated leukocytes was increased by 40% compared to the control, and the expression was decreased by 30% upon zerumbone treatment (Fig. 5A). Similarly, the ICAM-1 expression in the LPS group was increased by 57% compared to the control group and was decreased by 54% after treating with zerumbone (Fig. 5B).

**Figure 5.**
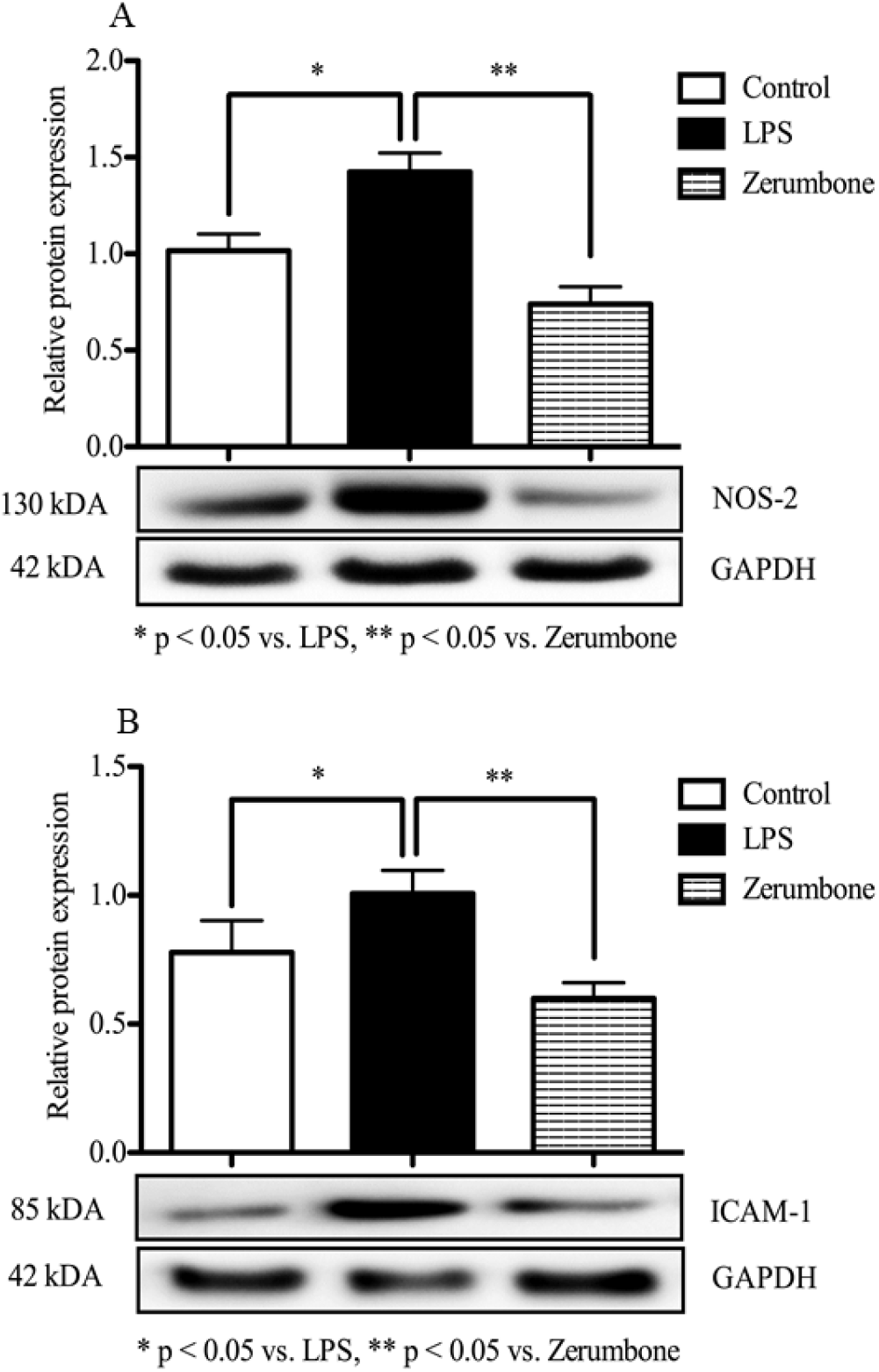
Protein expressions of NOS-2 (A), and ICAM-1 (B) in peripheral blood leukocytes. Values are Mean ± SD of 3 samples. A p value < 0.05 was considered statistically significant.

## Discussion

Phytomolecules with potential anti-eicosanoid properties offer a safe strategy to downregulate inflammation. This study demonstrated the anti-inflammatory effects of zerumbone by assessing its ability to intervene the eicosanoid pathway in peripheral blood leukocytes activated by lipopolysaccharides. Various investigators have explored Zerumbone to treat a wide range of complications [1, 26]. It has been shown that zerumbone when administered orally to mice, with doses up to 2000 mg/kg body weight, caused no significant changes in the circulatory cells and bone marrow [27]. Acute toxicity study of zerumbone-loaded nanostructured lipid carrier on BALB/c mice indicated the safety of zerumbone upon oral administration [28]. In this study, we explored the anti-eicosanoid properties of zerumbone, as eicosanoids have been implicated in the manifestation of pain and fever, consequently promoting full-blown inflammation [29]. We targeted leukocytes to assess the anti-eicosanoid effects of zerumbone, as leukocytes are known to initiate and propagate inflammatory reactions in many tissues, including the brain [30, 31]. The evidence from this study indicated that zerumbone positively interact with mPGES-1 (responsible for mitochondrial prostaglandins synthase) > NOS-2 (responsible for NO production) > FLAP (responsible for activation of 5-LOX) > COX-2 (responsible for PG synthesis) > LTA4-hydrolase (responsible for LTA4 conversion to LTB_4_) > BLT-1 (responsible for LTB_4_ induced chemotaxis) > EP-4 (responsible for PGE_2_ induced activation) > ICAM-1 (responsible for leukocyte docking to endothelium), in the order specified. The greater binding affinity of zerumbone with mPGES-1, NOS-2, FLAP, and COX-2 indicates the ability of zerumbone to intervene, nitric oxide, leukotriene, and prostaglandin generation, respectively. The expression of key enzymes and receptors of eicosanoid pathways were also measured to support the molecular docking observations. Decreased expression of NOS-2, 5-LOX, and COX-2 in leukocytes treated with zerumbone further supports the docking study results. Though zerumbone showed a greater binding affinity for FLAP, it did not alter the FLAP expression level (data not shown). We speculate that the interaction between zerumbone and FLAP lowers the activation of 5-LOX needed for leukotriene generation [32]. Besides, zerumbone mediated dampening of COX-2 enzyme activity and its interaction with the major receptor of PGE2 (EP-4) and LTB4 (BLT-1) may further weaken the eicosanoid signaling in leukocytes. As evident in this study, the LPS triggered production of proinflammatory lipid mediators, including PGE2, LTB4, and LTC4, were found to be diminished in peripheral blood leukocytes pre-treated with zerumbone, further underscores the inhibitory actions on the cyclooxygenases and lipoxygenases. Oxidative stress and inflammation are interlinked. Our study disclosed that zerumbone effectively prevents the generation of ROS induced by LPS in leukocytes, consequently weakening the enzymes and receptors of eicosanoid synthesis [33].

The downregulation of ICAM-1, an endothelial and leukocyte associated transmembrane protein that facilitates leukocyte-endothelial transmigration, may lower the oxidative and inflammatory trigger in the tissues and thus protect them from injury. The decreased ICAM-1 expression in leukocytes treated with zerumbone indicates that zerumbone treatment minimizes the leukocyte docking to the endothelium, a critical event in the inflammation triggered tissue injury [34]. Since studies have linked the PGE_2_/EP-1 signaling pathway for the increased expression of ICAM-1 and consequent inflammatory response, further studies on the eicosanoid mediated regulation of adhesion molecules need to be elucidated [35]. Besides, the zerumbone treatment also lowered the LPS induced nitric oxide production, possibly through the decreased expression of NOS-2, as evident from this study. Studies have shown that nitric oxide (NO) derived from NOS-2 modulate the biological levels of arachidonic acid-derived cell signalling molecules by altering the activity of cyclooxygenases due to Tyr nitration [36]. 3-Nitrotyrosine formation in proteins is considered a hallmark reaction of peroxynitrite, which can form via NOsuperoxide reactions in an inflammatory setting. Further, the reduction in the OS markers triggered by LPS in leukocytes by zerumbone underscores the ameliorating effect of zerumbone on leukocyte-mediated inflammation. In agreement with the above observation, the leukocyte’s well-kept ability to fight LPS induced free radicals can be justified by their restored antioxidant defense enzymes (catalase, SOD, glutathione peroxidase, and glutathione transferase). The overall anti-eicosanoid properties of zerumbone are graphically represented in Fig. 6. Therefore, this study provides first evidence for the anti-inflammatory properties of zerumbone via the downregulation of eicosanoids in peripheral blood leukocytes activated with lipopolysaccharides. We conclude that zerumbone, a monocyclic sesquiterpene present in the Zingiberaceae family, is a potential anti-eicosanoid molecule that needs to be explored further in the management of inflammatory complications.

**Figure 6.**
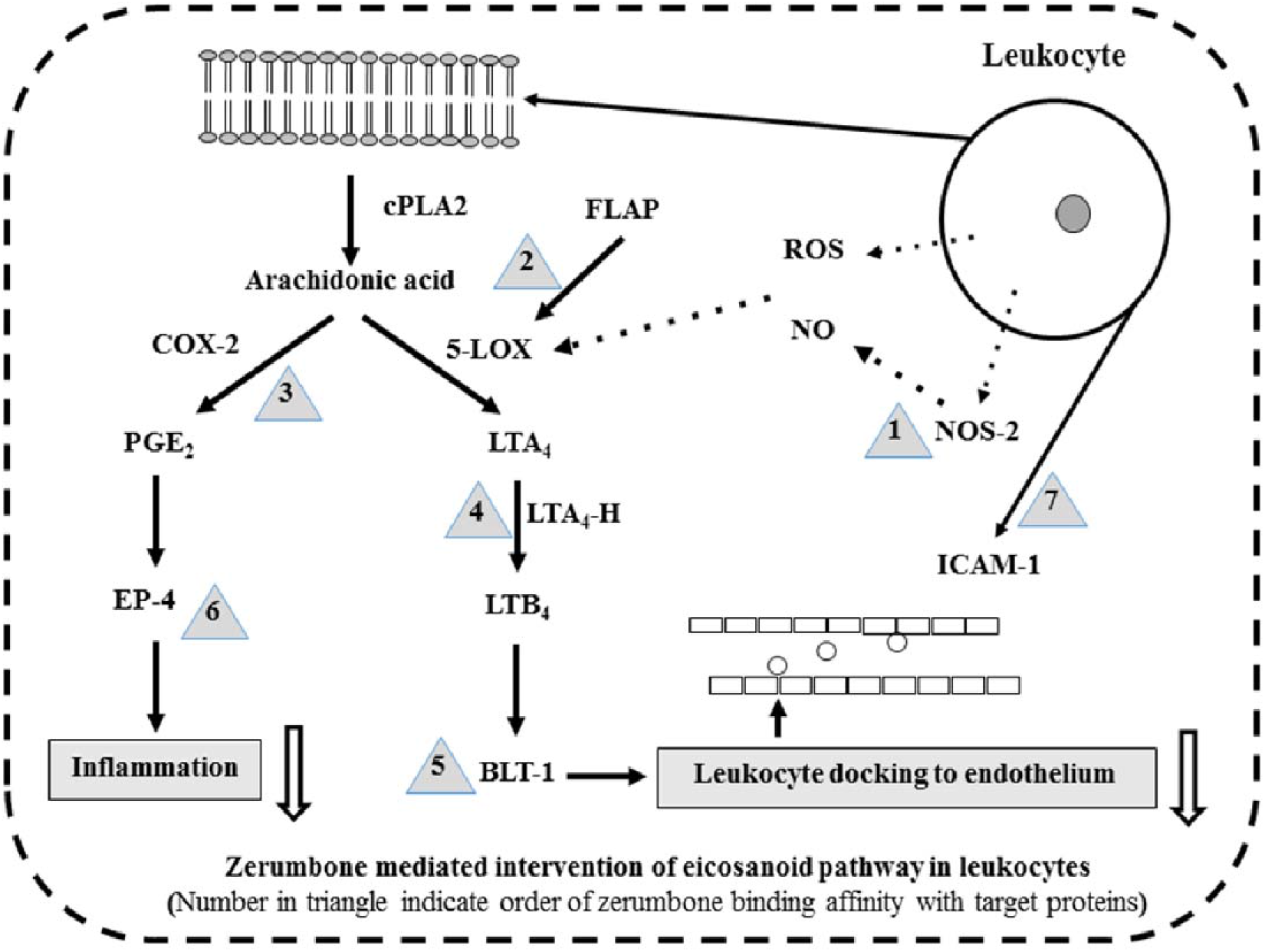
Graphical representation of zerumbone mediated intervention of eicosanoid pathway in leukocytes.

## Acknowledgments

Mr. Vinayak Uppin acknowledges DBT, New Delhi, for the Research Fellowship and CSIR-Central Food Technological Research Institute, Mysore, India, for providing the research facilities.

## ETHICS DECLARATION

### Credit Author Statement

Vinayak and Hamsavi were responsible for the execution of the experiment and VU for the data assessment. BBK was responsible for the zerumbone extraction. RRT was responsible for the research question, design, data assessment, and manuscript writing. All the authors contributed to this manuscript and approved the final version.

## Abbreviations

ICAM-1: inter cellular adhesion molecule-1
OS: oxidative stress
COX-2: cyclooxygenase-2
5-LOX: 5-lipoxygenase
FLAP: 5-lipoxygenase activating protein
mPGES-1: microsomal prostaglandin E synthase-1
NOS-2: nitric oxide synthase-2
ROS: reactive oxygen species
cPLA2: cytosolic phospholipase-A2
BLT-1: leukotriene B_4_ receptor-1
EP-4: prostaglandin-E receptor-4

