## Supplementary data for "Modulatory potentials of zerumbone isolated from ginger (*Zingiber zerumbet*) on eicosanoids: evidence from LPS induced peripheral blood leukocytes"

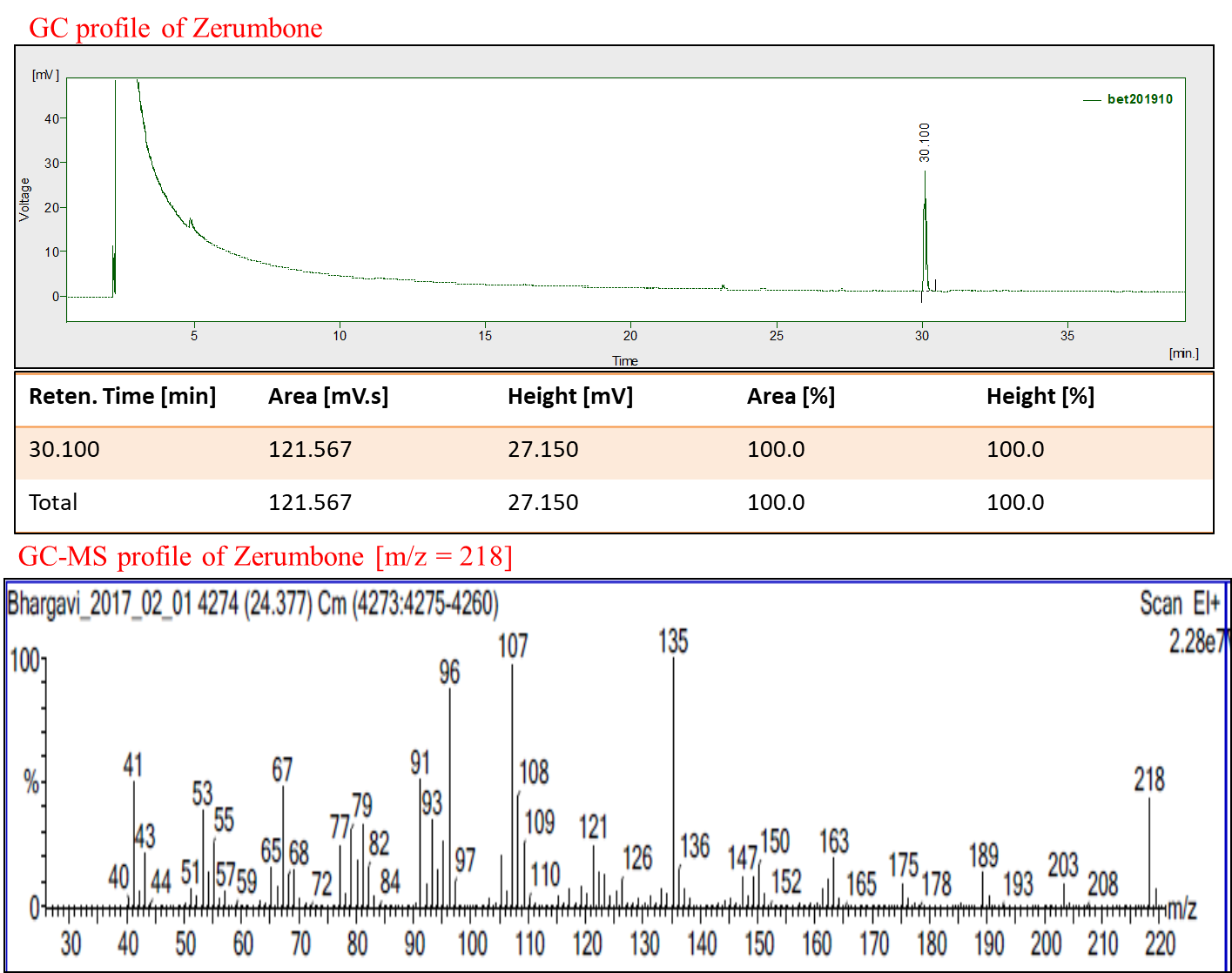

Figure 1 The GC and GC-MS chromatogram of the zerumbone crystals.

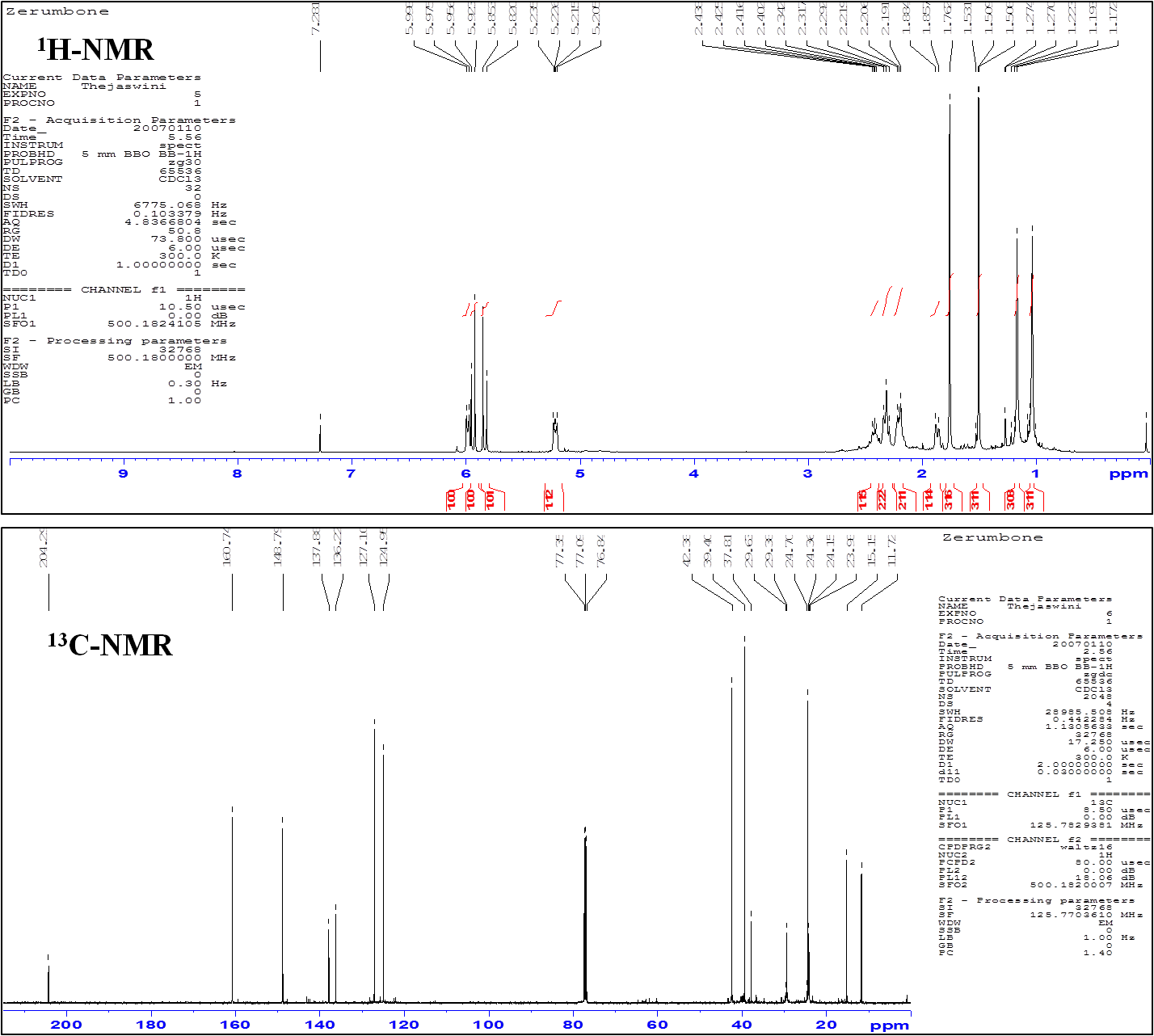

Figure 2 ^1^H & ^13^C NMR spectra of pure zerumbone crystals. The following are the NMR spectral and physical characteristics, which align with the literature values. Colorless crystalline solid; m.p. 64-66 °C; ^1^H NMR (500MHz, CDCl3): δ = 6.03 (d, 1H, J = 11.3 Hz ), 6.00 (d, 1H, J = 16.4 Hz), 5.88 (d, 1H, J = 16.4 Hz), 5.27 (d, 1H, J = 15.4 Hz), 2.43-2.50 (m, 1H, CH_2_), 2.36 (t, 2H, CH_2_ , J = 12.5 Hz), 2.20-2.30 (m, 2H, CH_2_), 1.89-1.94 (m, 1H, CH2), 1.81 (s, 3H, CH_3_), 1.56 (s, 3H, CH_3_), 1.22 (s, 3H, CH_3_) 1.08 (s, 3H, CH_3_); ^13^C NMR (125 MHz, CDCl_3_): δ = 160.37, 148.43, 137.66, 135.95, 126.87, 124.70, 42.12, 39.14, 37.54, 29.11, 24.08, 23.87, 14.88, 11.45.
